# AGR2, a potential prognostic and predictive biomarker and therapeutic to overcome drug resistance in HER2-overexpressing cancer

**DOI:** 10.1101/2025.07.23.666253

**Authors:** Sarai Martinez-Pacheco, Joaquín J Maqueda, Carla Oliveira, Lorraine O’Driscoll

## Abstract

The development of resistance to HER2-targeted drugs is common. Neratinib, an orally delivered cost effective drug is showing great promise for patients with HER2-overexpressing tumours but, unfortunately, for many the development of resistance is inevitable. In this study spanning novel pairs of cell line models and data from patients’ samples, we used a broad range of methodologies including mass-spectrometry-based quantitative proteomics, ectopic gene expression, RT-qPCR, immunoblotting, cytotoxicity, proliferation, invasion, and migration assays. We found loss of anterior gradient protein 2 (AGR2) to be prognostic for poor overall survival for patients with HER2-overexpressing tumours, in contrast to what has been reported for estrogen-positive breast cancer. Additionally, we discovered that loss of AGR2 is not only associated with neratinb-resistance, but that its restoration by ectopic AGR2 expression in neratinib-resistance cells partly restores their drug sensitivity. Furthermore, we established that keys metastatic phenotypic traits acquired with neratinib-resistance were blocked by ectopic AGR2 expression in the cells. Taken together, our results show that AGR2 has potential as a prognostic biomarker for outcome from HER2-overexpressing cancer; a predictive biomarker for resistance/response to neratinib; and a potential therapeutic protein for helping restore sensitivity to neratinib and overcoming aggressive cellular traits associated with cancer metastasis.

**Statement of Significance:** AGR2 loss in HER2-overexpressing cancer is prognostic for poor outcome and predictive for HER2-targeted drug-resistance, with AGR2 re-introduction helping restore drug sensitivity and overcoming phenotypic characteristics key to metastasis.

## Background

Breast cancer is reportedly the main cause of morbidity and mortality in women throughout the world^1^. Due to the development of HER2-targeted drugs, many of the approximately 20% of breast cancer patients with HER2-overexpressing tumours -who otherwise would have a poor prognosis-do well. However, it has been estimated that up to 70% of these patients do not gain meaningful long-term benefit, as a result of innate or acquired resistance/cross-resistance to HER2-targeted therapies^2^. Indeed, drug resistance is the main cause of these drugs failing in the clinic.

Neratinib is an orally delivered, irreversible HER2, EGFR and HER4 inhibitor^3^ offering substantial promise to patients. Neratinib was approved by the FDA and the EMA in 2017 and 2018, respectively, for extended adjuvant treatment of early stage HER2+ breast cancer. A phase I/II study (NCT00398567) showed neratinib to be effective for HER2+ breast cancer patients with acquired and innate resistance to trastuzumab. Additionally, its combination with trastuzumab was safe and well-tolerated, improving clinical outcomes in some subsets of patients^4^. A phase III trial (extended adjuvant treatment of breast cancer with neratinib, ExteNET) demonstrated a significant improvement on 5-year invasive disease-free survival in early-stage breast cancer patients, following trastuzumab-based adjuvant therapy (NCT00878709)^5^. Furthermore, an open-label phase II monotherapy trial is underway to establish how well neratinib performs in treating breast cancer that has spread to the brain (NCT01494662)^6^. Other recent studies have concluded that neratinib -as an extended adjuvant-is a feasible option after various anti-HER2 pre-treatments and its tolerability, mainly diarrhea, can be managed^1,7^. Results from the Phase III NALA trial have shown that for patients with HER2+ metastatic breast cancer, neratinib plus capecitabine significantly improved progression-free survival and time to intervention for CNS disease, when compared to lapatinib and capecitabine^8^. Furthermore, adding neratinib with capecitabine is more cost-effective than lapatinib with capecitabine^9^.

Although such evidence indicates that neratinib is a very promising drug even in comparison to other HER2-targeted drugs, resistance to it may occur and undermine its benefit. For that reason, understanding the ability of breast cancer cells to acquire neratinib resistance (and other associated undesirable effects on cancer cell behaviour) is necessary to move towards preventing or circumventing this resistance. Previously we established novel neratinib-resistant cell variants of HER2-positive breast cancer cell models and found that the development of resistance to neratinib confers cross-resistance to other HER2-targeted therapies including trastuzumab, lapatinib and afatinib, and that the resistant variants have a more aggressive phenotype compared to their drug-sensitive counterparts^10^; as often occurs in drug-resistant tumours in patients. However, the mechanism responsible for this remained unknown. Here, in studies spanning cell lines and data from patients’ samples, we aimed to investigate the mechanism responsible for neratinib-resistance and other associated phenotypic changes and to investigate a way towards overcoming the same.

## Materials and Methods

### Cell Culture

Eight HER2-positive breast cancer cell lines/variants HCC1954 and SKBR3, their respective neratinib-resistant (-NR) variants HCC1954-NR and SKBR3-NR^10,11^, as well as other variants established during the course of this study including HCC1954 NR-pcEMPTY, HCC1954-pcAGR2, SKBR3 NR-pcEMPTY, SKBR3-pcAGR2, were routinely maintained in RPMI-1640 medium (Sigma-Aldrich, Burlington, MA, USA; Cat. #: R0883) supplemented with 10% foetal bovine serum (FBS) (Thermo Fisher Scientific, Waltham, MA USA; Cat. #: 10270-106), and 2 mM L-Glutamine (Sigma-Aldrich, Cat. #: G7513) as complete medium. For studies involving EVs collection, cells were cultured in medium containing FBS that had been depleted of its EVs. All cells were cultured at 37 °C with 5% CO_2_ and routinely tested to ensure that they were free of *Mycoplasma* contamination. The authentication of all cell lines and variants were confirmed using STR profiling. All experiments described were performed a minimum of n=3 times.

### Global proteome profiling of drug-resistant versus drug-sensitive cells

To identify changes that may be associated with -and potentially causally involved in-drug-resistance, mass spectrometry-based quantitative proteomics with LC-MS/MS was used to analyse the protein expression profile (DEP) of HCC1954, HCC1954-NR, SKBR3 and SKBR3-NR cells. Differential protein expression analysis of the drug-resistant versus their drug-sensitive counterparts was investigated.

For protein extraction and processing for proteomics, each sample (1×10^6^ cells) was suspended in 6M urea (Sigma, Cat. #: U5378) and 50mM ammonium bicarbonate (ABC) (Honeywell, Cat. #: 1066-33-7). Samples were vortexed and incubated for 30 min at RT, sonicated on ice 20 sec (pulse time 5 sec, pulse off time 5 sec), and centrifuged at 16,000g for 10 min. The protein concentration of all the samples were determined by Bradford assay^12^ using Bio-Rad Protein Assay Dye Reagent (Bio-Rad, Cat. #: 500-0006). Samples were prepared at 50 μg protein in 30 μl solution for subsequent digestion.

For protein digestion by in-solution tryptic digestion, freshly prepared 100 mM dithiothreitol (DTT) (Sigma, Cat. #: D0632-1G) was added to the cell lysate samples using a ratio of sample volume/20 calculated for each sample to achieve a final concentration of 5 mM. Tubes were vortexed and incubated at 60°C for 30 min to disrupt disulphide bonds in proteins. Samples were centrifuged briefly to collect protein samples down in the tubes and 200 mM of iodoacetamide solution (IAA) (Sigma-Aldrich, Cat. #: I1149-5G) was added using a ratio of total sample volume (including DTT)/20, calculated independently for each sample. Samples were vortexed, incubated in the dark at RT for 30 min. and then diluted sufficiently with 50 mM ABC to reduce the concentration of urea to <2 M. A vial of trypsin singles (Sigma Aldrich, Cat. #: T7575) was added to each reduced alkylated sample which was then vortexed and incubated overnight at 37°C on a thermomixer (Eppendorf) at 350rpm. Trypsin digestion was stopped by adding HPLC grade acetic acid (Fisher Scientific, Cat. #: A/0406/PB08), 1% of the final volume, and samples were prepared for peptide desalting step by centrifugation at 10,000rpm for 5 min.

In preparation for quantitative LC-MS/MS analysis, the peptide samples were desalted using C18 ZipTip® pipette tips (Millipore, Cat. #: ZTC18S960) following the manufacturer’s protocol. Then, subjected to an evaporation process for 10-15 min using CentriVap concentrator, and resuspended in 25 μl of 0.5% acetic acid containing 2.5% ACN. Finally, they were centrifuged at 15,000 rpm at 4°C for 5 min and 20 μl supernatant transferred to the MS vial. To determine the peptide concentration of the samples, Pierce^TM^ Quantitative Colorimetric Peptide Assay (Thermo Scientific, Cat. #: 23275) was performed following the manufacturer’s instructions. Absorbance was read at 480 nm using a FluoStar Optima microplate reader (BMG Labtech). Samples were diluted to a final concentration of 40 ng/μl for the identification and quantification of the proteins using timsTOF Pro Mass Spectrometer (Bruker Daltonics, Bremen, Germany) coupled to a nanoElute (Bruker Daltonics, Bremen, Germany) ultra-high pressure nanoflow liquid chromatography system (UHPnLC). For further details see Supplemental Materials and Methods).

Protein identification involved raw data from the Bruker Timstof being processed using MaxQuant version 1.6.10.43, incorporating the Andromeda search engine^13^. To identify peptides and proteins, MS/MS spectra were matched to the Uniprot homo sapiens database (2020_05) containing 75,074 entries. All searches were performed with tryptic specificity allowing two missed cleavages. The database searches were performed with carbamidomethyl (C) as fixed modification and acetylation (protein N terminus) and oxidation (M) as variable modifications. Mass spectra were searched using the default setting of MaxQuant namely a false discovery rate of 1% on the peptide and protein level. For the generation of label free quantitative (LFQ) ion intensities for protein profiles, signals of corresponding peptides in different nano-HPLC MS/MS runs were matched by MaxQuant in a maximum time window of 1 min^14^.

In relation to data mining and bioinformatics analysis, the DEP package in R software version 1.4.1106 for Windows^15^ was used to analyse a total of 12 samples (*n* = 3, of each of the four cell variants). Protein hits identified from the reversed database were removed and protein groups were filtered for q-value < 1%. Label-free quantification (LFQ) intensities extracted by MaxQuant were log2-transformed and normalised by variance stabilisation. Only those proteins with LFQ intensity >0 and present in a minimum of two out of three replicates of each cell line variant were considered. Differential enrichment analysis was performed by applying empirical Bayes methods on protein-wise linear models using *limma* package^16^. False Discovery Rate (FDR) estimates were obtained by *fdrtool.* DEP were those with |log_2_(Fc)| > log_2_(1.5) and q-values (tail area-based FDR) <0.05. Gene Ontology and pathway analysis were performed by *enrichr* web tool^17^.

### Potential clinical relevance of AGR2 in HER2-overexpressing breast cancer

Before progressing to further investigate AGR2 i.e. the lead protein identified by proteomics, we wished to establish if this lead protein was likely to have clinical relevance. For this, association between AGR2 expression and overall survival (OS) was investigated for n=379 HER2-overexpressing breast tumours, using KM Plotter database (http://kmplot.com/analysis)^18^. Hazard ratios (HRs) with 95% confidence intervals and logrank P-value were determined.

### Stable transfection of AGR2 cDNA into drug-resistant cells

To perform AGR2 cDNA transfections, HCC1954 NR and SKBR3 NR cells were seeded at 2.5×10^5^ and 3.6×10^5^ cells/well, respectively, in six-wells plates (Costar, Cat. #: 3516) and cultured for 24 hrs. Plasmids containing AGR2 cDNA (EX-I0122-M94), EGFP cDNA (EX-EGFP-M94) and empty vector (EX-NEG-M94) were sourced from Genecopoeia. Following procedural optimisation, 2 µg of plasmids were transfected using FuGENE 4K (Promega, Cat. #: E5911) according to manufacturer’s protocol. Successful transfection was verified by fluorescence microscopy detecting expression of the integrated selection marker GFP. Commencing the following day, GFP-positive cells were selected in complete medium containing 2 µg/ml puromycin (Sigma-Aldrich; Cat. #: P4512). These cells were expanded as pools of puromycin-resistant cells after 4 weeks under selection pressure. Along with cells containing the empty vector (EX-NEG-M94), untransfected cells were used as an additional negative control. Stable populations of HCC1954 NR-pcAGR2, HCC1954 NR-pcEGFP, HCC1954 NR-pcEmpty, SKBR3 NR-pcAGR2, SKBR3 NR-pcEGFP, and SKBR3 NR-pcEmpty were established.

### RT-qPCR

cDNA from all 8 cell line variants was prepared using the SuperScript IV CellsDirect cDNA synthesis kit (Invitrogen, Cat. #: 11750150). Quantitative PCR (qPCR) was performed using PowerUP^TM^ SYBR^TM^ Green Master Mix (Invitrogen, Cat. #: A25741). Transfection efficiency was established by comparing AGR2 amplification normalised to the housekeeping gene β-actin in each sample. Primers: AGR2 primer (GeneCopoeia; Cat. #: HQP090714); KiCqStart^TM^ ACTB primers (Sigma-Aldrich; Cat. #: H_ACTB_1, NM_001101) were used. Amplification was performed in triplicate in a ViiA™ 7 Real-Time PCR System with 96-Well Block (Applied Biosystems™, Cat: # 4453534). The specificities of the products were confirmed by dissociation curve analysis.

### Immunoblotting

A total of 30 μg protein per sample was resolved on a 10% (Bio-Rad Laboratories, Cat. #: 4561036) or 7.5% Mini-PROTEAN TGX^TM^ 15-well gel (Bio-Rad Laboratories; Cat. #: 4561026), as appropriate, and the proteins were transferred to PVDF membranes (Bio-Rad Laboratories; Cat. #1620177). Membranes were then blocked with 5% (w/v) BSA in PBS containing 0.1% Tween-20 (PBS-T). Primary antibodies, incubated overnight at 4°C, were used in the following dilutions: anti-human AGR2 (1:1000, Abcam; Cat. #: ab76473), anti-HER2, (1:1000, Calbiochem; Cat. #: OP15), anti-E-Cadherin (1:1000, Abcam; Cat. #: ab40772), anti-N-Cadherin (1:1000, Abcam; Cat. #: ab76011), anti-Vinculin (1:1000, Cell Signaling; Cat. #: 13901S), anti-TGF-β1 (1:1000, Abcam; Cat. #: ab179695), and anti-β-actin (Sigma-Aldrich; Cat. #: A1978). After washing three times with PBS-T, the membrane was incubated with anti-Rabbit IgG (1:1000, Cell Signaling; Cat. #:7074) or anti-Mouse IgG (1:1000, Cell Signaling; Cat. #: 7076) in 3% BSA/PBS-T for 1 hr at room temperature (RT) and imaging was performed using an automated Chemidoc exposure system (Bio-Rad Laboratories). SuperSignal West Femto Chemiluminescent Substrate (Thermo Fisher Scientific; Cat. # 11859290) was used for detection. Densitometric analysis was performed using Fiji software^19^.

### Proliferation assays

HCC1954 and SKBR3 cell variants were seeded in 96-well plates (Costar, Cat. #: 3595) at a density of 3×10^3^ and 5×10^3^, respectively. Proliferation was measured after 24, 48, 72 and 96 hrs, using acid phosphatase assay method^20^.

### Neratinib’s cytotoxicity

Neratinib’s IC_50_ on HCC1954 and SKBR3 variants was determined using cytotoxic assay. Cells were seeded as for proliferations and 24 hrs later were exposed to increasing concentrations of neratinib ranging from 0–100 nM for HCC1954 and SKBR3; 0–900 nM HCC1954 NR cell variants; and 0–300 nM SKBR3 NR cell variants. After 5 days, cell viability was evaluated using the acid phosphatase method.

### Invasion assay

Invasion assays were performed using 8-μm pore size 24-well transwell chambers (Falcon, Cat. #: 353097). Inserts were pre-coated with extracellular matrix (ECM) (Sigma-Aldrich, Cat. #: E1270). HCC1954 variants (1×10^5^/insert) and SKBR3 variants (1×10^6^/insert) were seeded in the upper compartment and allowed to migrate for 48 hrs. Cells in the upper chamber were removed and invaded cells were stained with crystal violet. Staining was solubilised in 10% acetic acid and read at 595 nm.

### Migration (wound-healing) assay

HCC1954 and SKBR3 cell variants were seeded at 2.5×10^5^ and 3×10^5^ cells/well, respectively, in 24-well plates (Costar, Cat. #: 3524) and cultured for 24 hrs to confluence. Medium was removed from wells and the monolayer of cells was scratched using a p200 tip. 1% FBS-supplemented RPMI medium was carefully placed in each well and the wound size was captured by phase contrast microscopy using ImageJ/Fiji® plugin Wound-healing_size_tool^21^ at 0, 24 and 48 hrs. 2-way ANOVA with Tukey post-test was performed to assess multiple comparison.

### Statistical analyses

Statistical analyses were performed using GraphPad Prism version 9.1.9 for macOS (GraphPad Software). Data are presented as means ± SEM. For experiments comparing differences, two sets of data were performed by Student’s t-test while two-way ANOVA was used for comparison between groups (transfected vs transfected cells). For RT-qPCR analysis, mRNA fold change was calculated according to the ΔΔC_T_ value. One-way ANOVA with Tukey post-test was performed to assess statistical significance. * *p* < 0.05; ** *p* < 0.01; *** *p* < 0.001; *p* < 0.0001.

## Results

### Loss is associated with resistance to the HER2-targeted therapeutic neratinib

To identify the potential mechanism of neratinib resistance, protein(s) differentially expressed in neratinib-resistant cell variants versus their neratinib-sensitive counterparts were investigated by global proteomics using quantitative LC-MS/MS. A total of eighty significant differentially expressed proteins were identified. Of those, 39 were differentially expressed between HCC1954-NR and HCC1954 variants, and 40 between SKBR3-NR and SKBR3 variants. Notably, only 1 DEP, i.e. anterior gradient protein 2 (AGR2), was shared between the HCC1954-NR versus HCC1954 variants and the SKBR3-NR versus SKBR3 variants. Specifically, expression of AGR2 was down-regulated in both HCC1954-NR and SKBR3-NR cell variants with a fold-change of 46.85 (*p* = 4.06×10^−7^) and 4.92 (*p* = 8.69 ×10^−4^) compared with their respective neratinib-sensitive counterparts (**Fig. 1A-C**). To validate this discovery, immunoblotting analysis was performed on independent protein samples of the same cell line variants. Of note, densitometric analysis (**Fig. 1D**) of the immunoblotting results (**Fig. 2B**) confirmed the proteomics data i.e., AGR2 expression was significantly reduced with neratinib resistance.

**Fig. 1.**
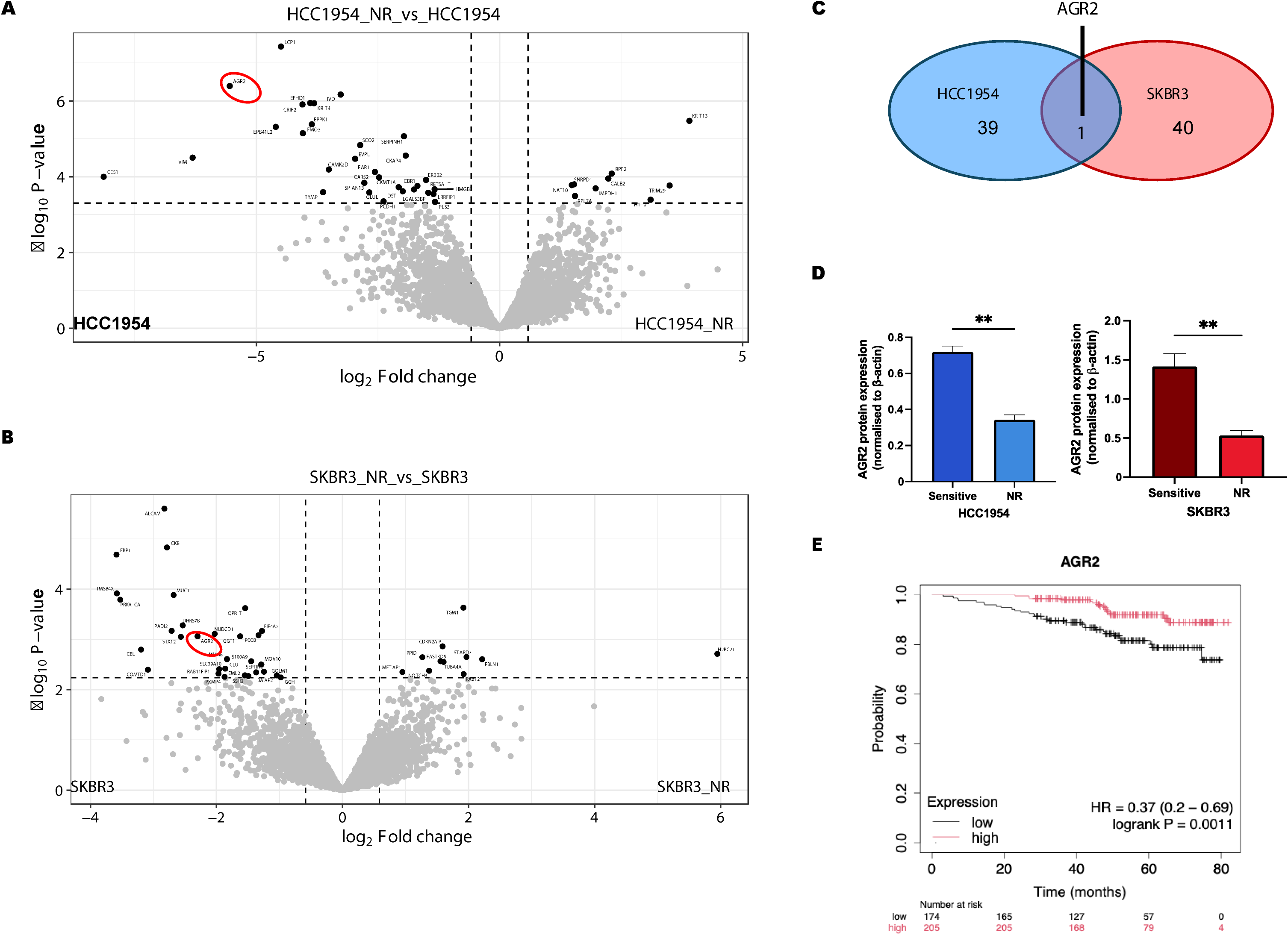
AGR2 in HER2 breast cancer is associated with overall survival, while its loss is associated with resistance to the HER2-targeted therapeutic neratinib. **(A)** Volcano plot for HCC1954 cell line variants, highlighting differentially expressed proteins; **(B)** Volcano plot for SKBR3 cell line variants, highlighting differentially expressed proteins. Dashed lines indicated fold-change and p-value cutoffs (|log_2_(Fc)| >= log_2_(1.5) and *p* <= 0.05, respectively) Par = neratinib-sensitive cell lines; NR = neratinib-resistant cell lines; HC = HCC1954; SK = SKBR3; **(C)** Venn diagram showing the differentially expressed proteins from comparative proteomic analysis and **(D)** validation of those findings by immunoblotting. **(E)** AGR2 is expressed in HER2-positive breast cancer, with high level expression significantly associated with favourable outcome in terms of overall survival and loss associated with poor outcome (HR = 0.37, *p* = 0.0011; duration = 60 months analysis, Kaplan-Meier graph represents *n* = 379 patients). ** *p* < 0.01.

**Fig. 2.**
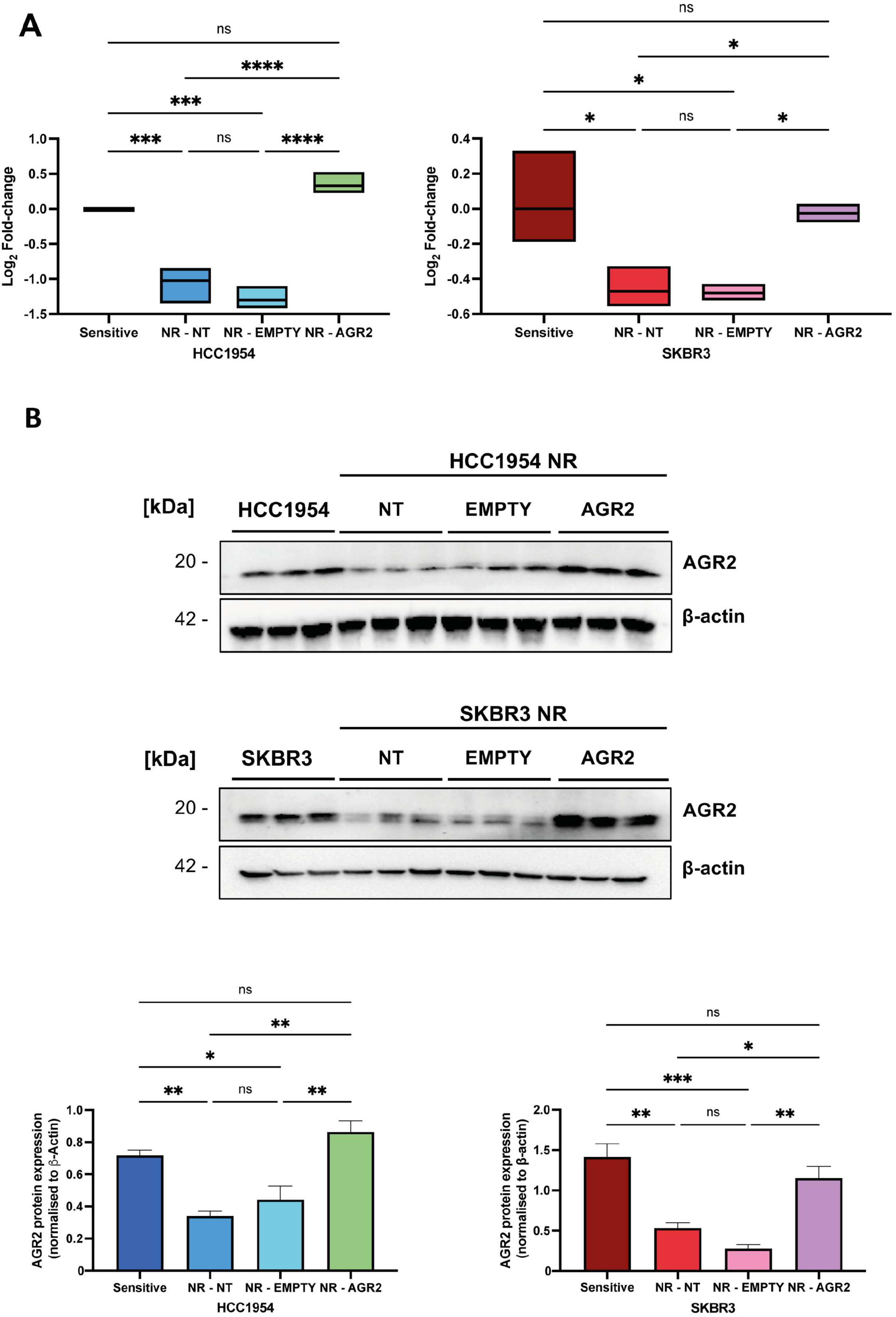
Loss of AGR2 mRNA and proteins with neratinib-resistance was restored to AGR2 levels in drug-sensitive cells by ectopically expressing AGR2. **(A)** AGR2 mRNA was analysed by RT-qPCR, with ACTB serving as an endogenous control for data normalisation. Amounts of AGR2 mRNA after AGR2 re-introduction (NR-AGR2), compared to those transfected with an empty plasmid (NR-empty) and with non-transfected (NR-NT) cells, were statistically significant for both cell line variants. (**B**) Similarly, expression of AGR2 protein, analysed by immunoblotting and its associated densitometric analysis normalised to !-actin, confirmed restoration of AGR2 protein. Data are represented as the mean ± SEM, n = 3. One-way ANOVA with Tukey post-test was performed for the multiple comparisons. ns = not significant; * *p* < 0.05; ** *p* < 0.01; *** *p* < 0.001; **** *p* < 0.0001.

### AGR2 in HER2 breast cancer is associated with survival, while its loss is prognostic for poor outcome

To determine whether AGR2 has relevance in outcome for patients with breast cancer, rather than being solely a cell line related observation, as a first step we mined microarray data relating to n=379 HER2-overexpressing breast tumours (given that -omics data is not yet available specifically related to human tumours resistant to neratinib). Kaplan-Meier estimates showed that the overall survival probability over time was significantly shorter (HR = 0.37 (0.35–0.56), *p* = 0.0011, **Fig. 1E**) for patients whose tumours had low AGR2 expression (*n* = 174) compared to those with higher expression (*n* = 205), suggesting AGR2’s potential as a cellular biomarker of resistance.

### AGR2 was successfully ectopically expressed in neratinib-resistant cell variants

Given that we found AGR2 to be significantly decreased in the neratinib-resistant cell lines compare to their drug-sensitive counterparts, and that low expression in breast tumours is associated with patients’ poor overall survival, we progressed to investigating if AGR2 likely represents a causal component of the mechanism of neratinib-resistance and, if so, if its re-introduction may be a therapeutic approach to overcoming resistance.

For this purpose, the resistant HCC1954-NR and SKBR3-NR cells were stably transfected with AGR2, EGFP (as marker of successful transfection), or EMPTY plasmid (as a control). Successful transfection and AGR2 overexpression were verified by microscopy after puromycin selection (images not shown), as well as by evaluating AGR2 expression by both RT-qPCR (**Fig. 2A**) and immunoblotting (**Fig. 2B**). Indeed, successful transfection with AGR2 cDNA significantly increased AGR2 mRNA transcription and protein translation in both HCC1954-NR and SKBR3-NR cells; achieving similar AGR2 levels of AGR2 mRNA and proteins to those in their respective neratinib-sensitive counterparts.

### Ectopic expression of AGR2 in neratinib-resistant cell variants does not affect proliferation, but decreases resistance/partially restores sensitivity to neratinib

A possible influence of AGR2 expression on cell growth, investigated over a 96 hrs time course (using acid phosphatase assay), showed no notable proliferative or anti-proliferative effect in either HCC1954 or SKBR3 cell variants (**Fig. S1**).

To establish if a potential direct relationship exists between AGR2 expression and resistance/sensitivity to neratinib, IC_50_ values of neratinib for HCC1954 and SKBR3 variants were subsequently determined (**Fig. 3)**. This analysis showed that although neratinib-resistant cell variants ectopically expressing AGR2 remained somewhat neratinib-resistance when compared with the age-matched neratinib-sensitive control cells, partial but significant re-sensitisation to neratinib (i.e., a decrease in the IC_50_) was observed in that comparison (and when compared with NR-pcEMPTY transfected cell variants). As expected, no significant differences in neratinib sensitivity were found between -NR and NR-pcEMPTY cells, demonstrating the reduction in resistance/partial restoration of sensitivity to neratinib was directly associated with increased AGR2 expression rather than due to simply transfecting/manipulating the cells.

**Fig. 3.**
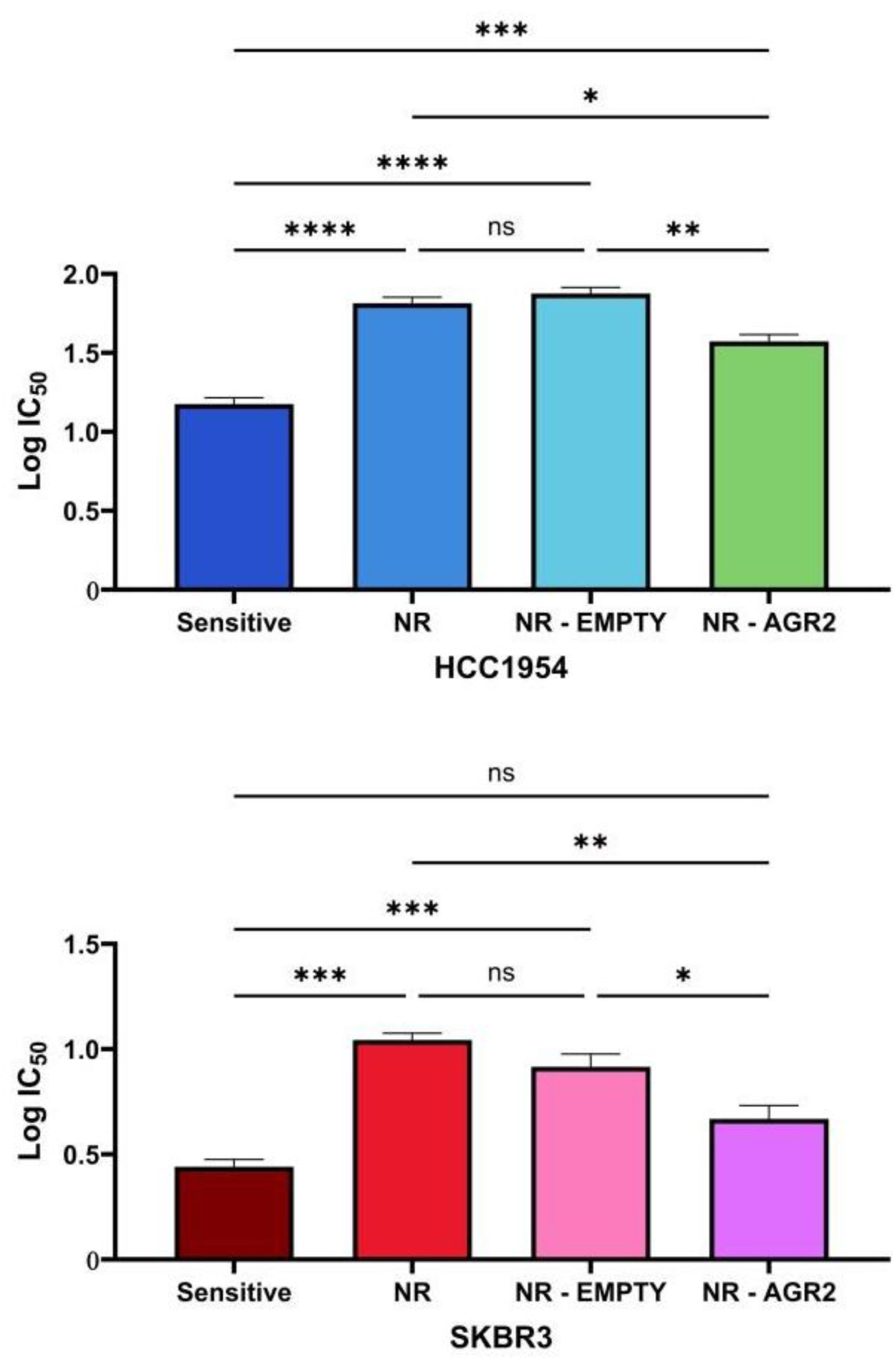
Ectopic expression of AGR2 in neratinib-resistant cell variants partially restore sensitivity to neratinib. The log (IC_50_) values for neratinib of (**A**) HCC1954 and (**B**) SKBR3 cell variants was determined using cytotoxic assays. After 4 days culture in the presence of neratinib partial, but significant, restoration of neratinib-sensitivity in both the previous drug-resistant variants was observed. The data represent the means from three independent experiments, with error bars showing the SEM of the means. One-way ANOVA was performed for the multiple comparisons. * *p* < 0.05; ** *p* < 0.01; *** *p* < 0.001; **** *p* < 0.0001.

### Ectopic AGR2 expression may prevent some key steps in the metastasis cascade

Metastasis is the spread of cancers throughout the body to where they are difficult to investigate and eradicate. For this, cancer cells must detach from the primary tumour, invade and move through adjacent tissues into blood or lymphatic vessels (an event known as intravasate) and, subsequently, invade out of the vessels and move into secondary sites (i.e. extravasate).

Invasion through ECM and migration of tumour cells through nearby tissues is crucial to entering blood vessels or lymphatic vessels (i.e., intravasation) and, again, for cells to move out of blood/lymphatic vessels (i.e., extravasation) and occupy the secondary/metastatic site^22^. In relation to invasion through ECM, neratinib-resistant HCC1954 cells (i.e., HCC1954-NR and HCC1954 NR-pcEMPTY variants) were significantly more invasive when compared to neratinib-sensitive HCC1954 cells. Interestingly, when HCC1954-NR cells were transfected with pcAGR2, invasion capacity reverted to that of drug-sensitive HCC1954 cells, i.e. no significant differences in invasion were found between HCC1954 NR-pcAGR2 and HCC1954 cells (**Fig. 4**).

**Fig. 4.**
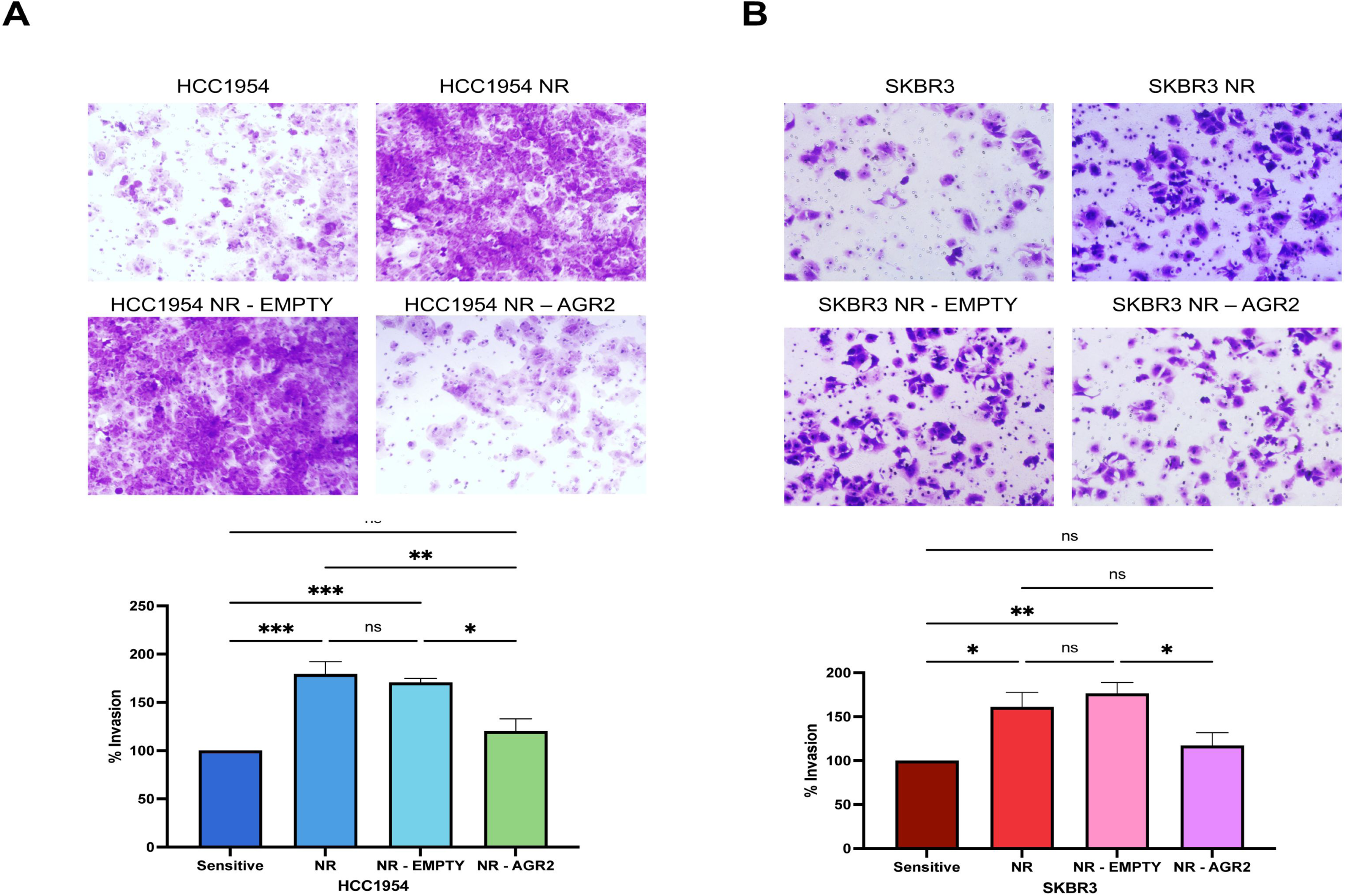
Ectopic expression of AGR2 in the significantly more invasive neratinib-resistant cells reduced cellular invasiveness. (**A**) Representative images of (**A**) HCC1954 and (**B**) SKBR3 invasive cells after 48 hrs seeding. Resistance to neratinib is significantly associated with increased cellular invasiveness through ECM. Ectopic expression of AGR2 reduced invasiveness of the neratinib-resistant cells to that of neratinib-sensitive cells. Images were taken at a 10X magnification; Graphs show the % of invasion at 48 hrs. The data are expressed as the mean ± SEM obtained from three independent experiments. 2-way ANOVA with Tukey post-test was performed for the multiple comparisons. * *p* < 0.05; ** *p* < 0.01; *** *p* < 0.001.

Similarly, when we investigated invasion in SKBR3 cell variants with ectopic AGR2 expression, we obtained a similar reduction in invasiveness in SKBR3-NR cell variant (**Fig. 4**). Indeed, no significant differences in invasiveness were found between neratinib-sensitive SKBR3 cells and SKBR3 NR-pcAGR2 cells, while SKBR3-NR and SKBR3 NR-pcEMPTY were more invasive compared to SKBR3 and SKBR3 NR-pcAGR2.

Using the wound-healing assay to evaluate migration, ectopic AGR2 expression in neratinib-resistant cells reduced migration to approximately that of neratinib-sensitive cells, for both HCC1954 and SKBR3 cell lines (**Fig. 5**). Specifically, when evaluated at both 24 hrs and 48 hrs time points, HCC1954 and NR-pcAGR2 had significantly decreased wound closure when compared to HCC1954-NR and HCC1954 NR-pcEMPTY cells. Similarly, SKBR3 and SKBR3 NR-pcAGR2 cells had significantly decreased wound closure when compared to SKBR3-NR and SKBR3 NR-pcEMPTY cells.

**Fig. 5.**
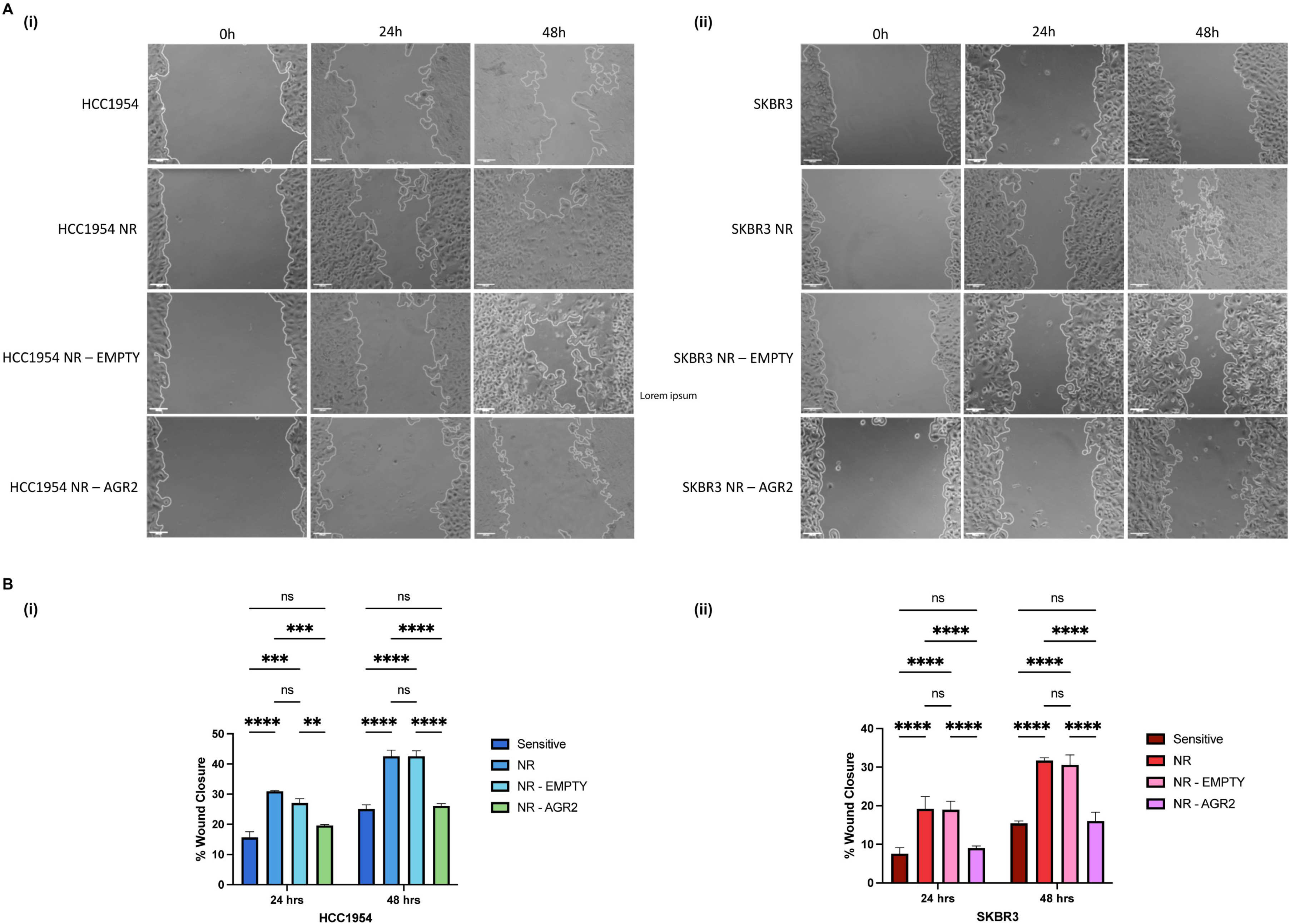
Ectopic expression of AGR2 in the significantly more migratory neratinib-resistant cells reduced cellular migration. (**A**) Representative wound-healing images of HCC1954 (**i**) and SKBR3 (**ii**) cell variants at 0, 24, and 48 hrs. Images were taken at a 10X magnification. Scale bar 100μm; (**B**) Graphs show the % of wound closure at 24 and 48 hrs in HCC1954 (**i**) and SKBR3 (**ii**) cell variants. Resistance to neratinib is significantly associated with increased cellular migration, while ectopic AGR2 expression reduced migration of the neratinib-resistant cells to that of neratinib-sensitive cells. Graph bars represent the means from three independent experiments, with error bars showing the SEM of the means. Graph bars represent the means from three independent experiments, with error bars showing the SEM of the means. Two-way ANOVA with Tukey post-test was performed for the multiple comparisons. * *p* < 0.05; ** *p* < 0.01; *** *p* < 0.001.

Together, neratinib resistant HCC1954-NR and SKBR3-NR variants were significantly more invasive and migratory than their drug-sensitive counterparts and ectopic AGR2 expression of AGR2 prevented these key aspects of the metastasis journey.

## Discussion

Anterior gradient 2 (AGR2), encoded by its gene on chromosome 7p21 in the human genome, is a 19kDa / 154 amino acid pro-oncogenic protein localised in the endoplasmic reticulum^23^. AGR2 was originally discovered in the African clawed frog *Xenopus laevis* where it plays an important role in embryonic development and. more recently, its importance in embryonic human development was also established^24^. AGR2 is frequently deregulated in cancer and has been implicated in many adverse events. A recent analysis of AGR2 protein using tissue microarrays representing 134 tumour categories, reported AGR2 positivity in 103/134 (77%) of tumour categories analysed^25^.

The relevance and potential causal role of AGR2 in breast cancer, and particularly HER2-positive breast cancer, has been largely unknown until now. AGR2 has been reported as present in estrogen receptor (ER) breast cancer cells lines, but not in ER-cell lines^15^, suggesting it to be of relevance in some breast cancer subtypes but not in others. An immunohistochemistry study of a tumour microarray (TMA) representing 315 tumours from patients with breast cancer showed AGR2 expression was prognostic for poor outcome, with only 26% of those with AGR2-positive primary tumours surviving to the 20 years follow up, compared to 96% of those with AGR2-negative tumours reaching this important timeline. Specifically, that study showed AGR2 expression to be significantly associated with estrogen receptor (ER) and progesterone receptor (PR) expression, and with cancer spread to the lymph nodes. However, no significant association was found with expression of HER2^26^. Another example study using breast tissue-specific TMAs again reported AGR2 expression to correlate significantly with expression of ER and PR -and also with androgen receptor (AR) and EGFR-but with no significant association found with HER2^27^. That study reported AGR2 expression as associated with nodal involvement and with poor outcome in response to hormone therapy. A subsequently study showed that AGR2 expression was not associated with either relapse-free survival (RFS) or overall survival (OS) when considering the whole breast cancer cohort evaluated (n=211), although when only those HER2-overexpressing (n=73) tumours were included, AGR2 expression was associated with RFS but not overall survival. Overall, evidence consistently associated AGR2 expression with ER/PR expression in breast cancer. Furthermore, the presence of AGR2 had been associated with resistance to tamoxifen and fulvestrant in ER-positive tumours^15,28^ and with resistance to the classical chemotherapy drug adriamycin/doxorubicin in breast cancer cell lines models, MCF-7 (ER^+^/PR^+/−^/HER2^−^) and MDA-MB-231 (ER^−^/PR^−^/HER2^−^)^29^. Other research suggests, however, that AGR2’s involvement in breast cancer is complex and that generalisation about AGR2 and breast cancer should not be made without evidence, given the maybe counterintuitive observation that low expression of AGR2 is associated with improved overall survival in breast cancer bone metastasis^30^.

As indicated, the data on AGR2 in relation to HER2-positive breast cancer is thus far conflicting and limited to its presence/absence, rather than potential causal and therapeutic roles. Addressing this, here we showed that AGR2 is, indeed, expressed by HER2-positive cell lines. This was established using a proteomics approach and validated by immunoblotting. Moving on to consider the potential clinical relevance of AGR2 in HER2 expressing tumours, we found that within this cohort, high expression of AGR2 was significantly prognostics for favorable outcome in terms of overall survival. Furthermore, our study made the discovery that AGR2 expression is significantly reduced in cells resistant to neratinib. Again, this was initially identified by proteomics and then validated by a second method, immunoblotting. This suggests that in HER2-positive breast cancer the presence of HER2 is favorable, and that loss of AGR2 may be predictive of resistance to HER2-targeting neratinib and poor outcome for patients.

Progressing to establish if AGR2 is likely to simply be associated with -or causally involved-in neratinib resistance, we then ectopically re-introduced AGR2 into neratinib resistant cells and, in so doing so, achieved partially restoration of neratinib sensitivity. This suggest that AGR2 is functionally involved in neratinib resistance and in restoration of sensitivity to the same.

In additional to development of drug resistance, cancer metastasis to secondary sites is a major reason for why patients die from cancer. Using laboratory methodologies to mimic major events contributing to cancer cell metastasis (i.e. cancer cell breaking from the primary tumour, invading though the extracellular matrix and moving towards blood or lymphatic vessels and similarly invading though ECM at the secondary site and moving to a place to colonise and grow), our findings indicate that ectopic AGR2 expression may also play a role in blocking these key undesirable events associated with metastasis.

Altogether, this study points to AGR2 as a favourable prognostic marker in HER2 breast cancer, with its loss potentially serving as a predictive biomarker for neratinib resistance in HER2-overexpressing cancers, and its administration as a therapeutic approach to blocking resistance and so adding value to HER2-targeting with neratinib.

## Authors Contributions

SM-P: designed and conducted experiments, analysed, and interpreted the data, and drafted the manuscript; JJ and CO: involved in the data analysis discussions, provided analytical tools, and edited manuscript; LOD: conceptualisation, formal analysis, funding acquisition, project administration, resources, supervision, writing - review & editing.

## Data availability statement

Supporting data can be found in the Supplementary Material. Otherwise, the data generated in this study are available upon request from the corresponding author.

## Supporting Information

### S1 Material and Methods

#### Quantitative LC-MS/MS analysis

The peptides were separated on a reversed-phase C18 Aurora column (25cm x 75μm ID, C18, 1.6 μm; IonOpticks, Australia) at a constant flow rate of 250 nL/min and an increasing acetonitrile gradient. Mobile phases were 0.1% (v/v) formic acid in water (phase A) and 0.1% (v/v) formic acid in acetonitrile (phase B). The peptides were separated by a gradient starting from 2% of mobile phase B and increased linearly to 32% for 60 min. This was stepped up to 95% of mobile phase B where it was maintained for 7 min. The injection volume was 5 μL, equivalent to a loading of 200 ng per sample.

The timsTOF Pro mass spectrometer was operated in positive ion polarity with TIMS (Trapped Ion Mobility Spectrometry), and PASEF (Parallel Accumulation Serial Fragmentation) modes enabled. The accumulation and ramp times for the TIMS were both set to 100 msec., with an ion mobility (1/k0) range from 0.62 to 1.46 Vs/cm. Spectra were recorded in the mass range from 100 to 1,700 m/z. The precursor MS Intensity Threshold was set to 2,500 and the precursor Target Intensity set to 20,000. Each PASEF cycle consisted of one MS ramp for precursor detection followed by 10 PASEF MS/MS ramps, with a total cycle time of 1.16 s.

## Supplemental Data sample

**Figure S1.**
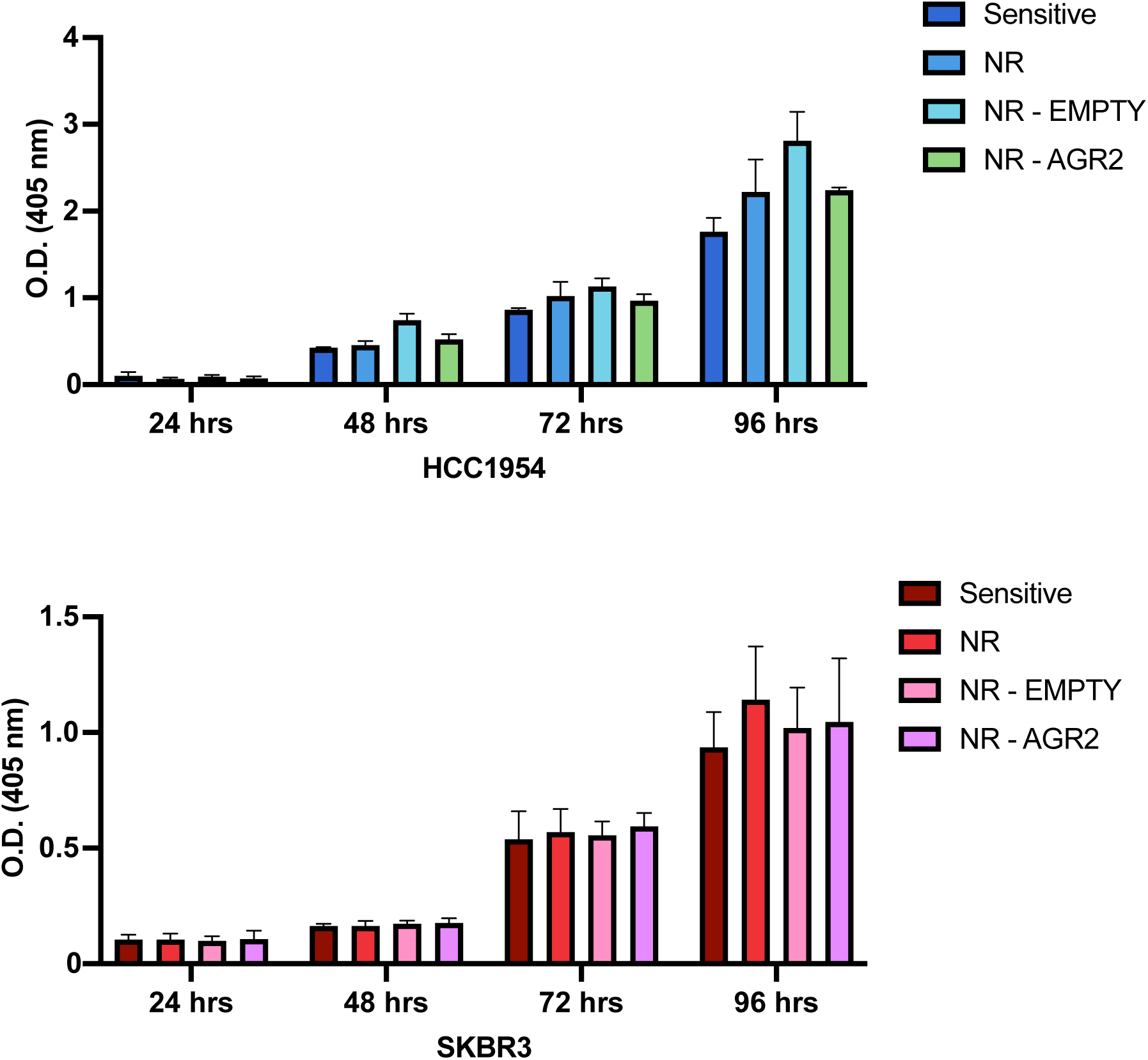
Ectopic expression of AGR2 does not affect proliferation rate of neratinib-resistant cell variants. Acid phosphatase assay results at 24, 48, 72, and 96 hrs. The data represent the means from three independent experiments, with error bars showing the SEM of the means. OD_405_, optical density at 405nm.

**Figure S2.**
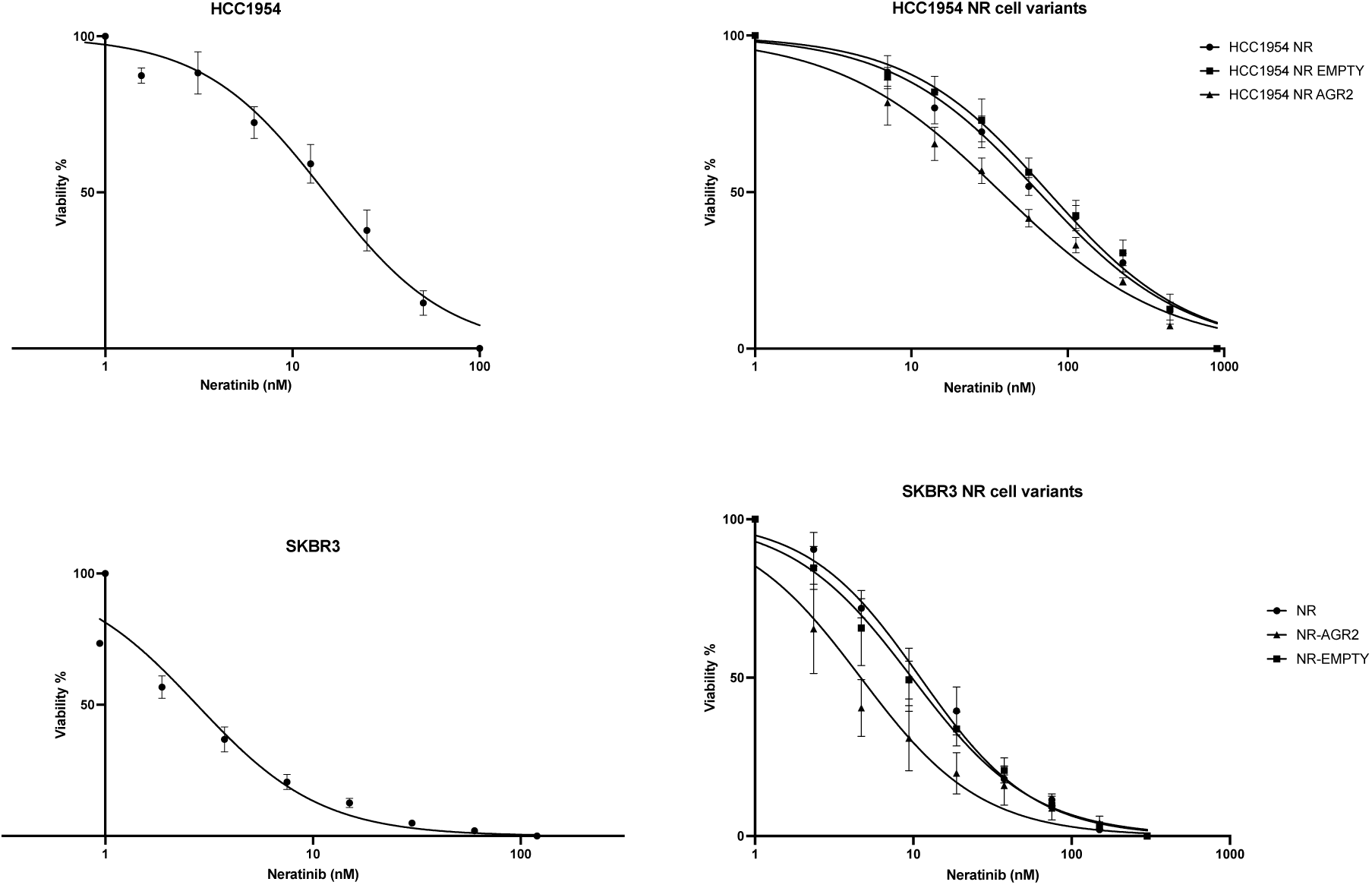
Drug sensitivity assay–dose response curves of HCC1954 and SKBR3 cell variants. Cytotoxicity assay on HCC1954 and SKBR3 cell variants after 5 days of neratinib treatment. IC_50_ curves represent *n*=3 biological repeats as mean ± SEM.

## References

1. Caputo R, Buono G, Lauro VD, et al. Neratinib as adjuvant therapy in patients with HER2 positive breast cancer: expert opinion. Future Oncol. 2023;19(24):1695–1708. doi:10.2217/fon-2023-0361

2. Vernieri C, Milano M, Brambilla M, et al. Resistance mechanisms to anti-HER2 therapies in HER2-positive breast cancer: Current knowledge, new research directions and therapeutic perspectives. Critical Reviews in Oncology/Hematology. 2019;139:53–66. doi:10.1016/j.critrevonc.2019.05.001

3. Bose P, Ozer H. Neratinib: an oral, irreversible dual EGFR/HER2 inhibitor for breast and non-small cell lung cancer. Expert Opinion on Investigational Drugs. 2009;18(11):1735–1751. doi:10.1517/13543780903305428

4. Blackwell KL, Zaman K, Qin S, et al. Neratinib in Combination With Trastuzumab for the Treatment of Patients With Advanced HER2-positive Breast Cancer: A Phase I/II Study. Clinical Breast Cancer. 2019;19(2):97–104.e4. doi:10.1016/j.clbc.2018.12.011

5. Delaloge S, Cella D, Ye Y, et al. Effects of neratinib on health-related quality of life in women with HER2-positive early-stage breast cancer: longitudinal analyses from the randomized phase III ExteNET trial. Ann Oncol. 2019;30(4):567–574. doi:10.1093/annonc/mdz016

6. Freedman RA, Gelman RS, Agar NYR, et al. Pre- and Postoperative Neratinib for HER2-Positive Breast Cancer Brain Metastases: Translational Breast Cancer Research Consortium 022. Clin Breast Cancer. 2020;20(2):145–151.e2. doi:10.1016/j.clbc.2019.07.011

7. Harbeck N, Wrobel D, Zaiss M, et al. Neratinib as Extended Adjuvant Treatment of HER2-Positive/HR-Positive Early Breast Cancer Patients in Germany, Austria, and Switzerland: Interim Results of the Prospective, Observational ELEANOR Study. Breast Care. 2024;19(1):1–9. doi:10.1159/000533657

8. Saura C, Oliveira M, Feng YH, et al. Neratinib Plus Capecitabine Versus Lapatinib Plus Capecitabine in HER2-Positive Metastatic Breast Cancer Previously Treated With ≥ 2 HER2-Directed Regimens: Phase III NALA Trial. J Clin Oncol. 2020;38(27):3138–3149. doi:10.1200/JCO.20.00147

9. Ren L, Ren N, Zheng Y, Yang Y, Xu Q. Economic evaluation of third-line neratinib plus capecitabine versus lapatinib plus capecitabine with HER2+ metastatic breast cancer. Front Oncol. 2023;13:1221969. doi:10.3389/fonc.2023.1221969

10. Breslin S, Lowry MC, O’Driscoll L. Neratinib resistance and cross-resistance to other HER2-targeted drugs due to increased activity of metabolism enzyme cytochrome P4503A4. Br J Cancer. 2017;116(5):620–625. doi:10.1038/bjc.2016.445

11. Martinez-Pacheco S, O’Driscoll L. Evidence for the Need to Evaluate More Than One Source of Extracellular Vesicles, Rather Than Single or Pooled Samples Only, When Comparing Extracellular Vesicles Separation Methods. Cancers. 2021;13(16):4021. doi:10.3390/cancers13164021

12. Bradford MM. A rapid and sensitive method for the quantitation of microgram quantities of protein utilizing the principle of protein-dye binding. Analytical Biochemistry. 1976;72(1-2):248–254. doi:10.1016/0003-2697(76)90527-3

13. Cox J, Neuhauser N, Michalski A, Scheltema RA, Olsen JV, Mann M. Andromeda: A Peptide Search Engine Integrated into the MaxQuant Environment. J Proteome Res. 2011;10(4):1794–1805. doi:10.1021/pr101065j

14. Cox J, Hein MY, Luber CA, Paron I, Nagaraj N, Mann M. Accurate Proteome-wide Label-free Quantification by Delayed Normalization and Maximal Peptide Ratio Extraction, Termed MaxLFQ. Molecular & Cellular Proteomics. 2014;13(9):2513–2526. doi:10.1074/mcp.M113.031591

15. Zhang X, Smits AH, van Tilburg GB, Ovaa H, Huber W, Vermeulen M. Proteome-wide identification of ubiquitin interactions using UbIA-MS. Nat Protoc. 2018;13(3):530–550. doi:10.1038/nprot.2017.147

16. Ritchie ME, Phipson B, Wu D, et al. limma powers differential expression analyses for RNA-sequencing and microarray studies. Nucleic Acids Research. 2015;43(7):e47–e47. doi:10.1093/nar/gkv007

17. Kuleshov MV, Jones MR, Rouillard AD, et al. Enrichr: a comprehensive gene set enrichment analysis web server 2016 update. Nucleic Acids Res. 2016;44(W1):W90–W97. doi:10.1093/nar/gkw377

18. Győrffy B. Survival analysis across the entire transcriptome identifies biomarkers with the highest prognostic power in breast cancer. Computational and Structural Biotechnology Journal. 2021;19:4101–4109. doi:10.1016/j.csbj.2021.07.014

19. Schindelin J, Arganda-Carreras I, Frise E, et al. Fiji: an open-source platform for biological-image analysis. Nat Methods. 2012;9(7):676–682. doi:10.1038/nmeth.2019

20. O’Driscoll L, Linehan R, Liang YH, Joyce H, Oglesby I, Clynes M. Galectin-3 expression alters adhesion, motility and invasion in a lung cell line (DLKP), in vitro. Anticancer Res. 2002;22(6A):3117–3125.

21. Suarez-Arnedo A, Torres Figueroa F, Clavijo C, Arbeláez P, Cruz JC, Muñoz-Camargo C. An image J plugin for the high throughput image analysis of in vitro scratch wound healing assays. Chirico G, ed. PLoS ONE. 2020;15(7):e0232565. doi:10.1371/journal.pone.0232565

22. Polacheck WJ, Zervantonakis IK, Kamm RD. Tumor cell migration in complex microenvironments. Cell Mol Life Sci. 2013;70(8):1335–1356. doi:10.1007/s00018-012-1115-1

23. Petek E, Windpassinger C, Egger H, Kroisel PM, Wagner K. Localization of the human anterior gradient-2 gene (AGR2) to chromosome band 7p21.3 by radiation hybrid mapping and fluorescencein situ hybridisation. Cytogenet Cell Genet. 2000;89(3-4):141–142. doi:10.1159/000015594

24. Jach D, Cheng Y, Prica F, Dumartin L, Crnogorac-Jurcevic T. From development to cancer - an ever-increasing role of AGR2. Am J Cancer Res. 2021;11(11):5249–5262.

25. Schraps N, Port JC, Menz A, et al. Prevalence and Significance of AGR2 Expression in Human Cancer. Cancer Med. 2024;13(21):e70407. doi:10.1002/cam4.70407

26. Barraclough DL, Platt-Higgins A, de Silva Rudland S, et al. The metastasis-associated anterior gradient 2 protein is correlated with poor survival of breast cancer patients. Am J Pathol. 2009;175(5):1848–1857. doi:10.2353/ajpath.2009.090246

27. Lacambra MD, Tsang JYS, Ni YB, Chan SK, Tan PH, Tse GM. Anterior Gradient 2 is a Poor Outcome Indicator in Luminal Breast Cancer. Ann Surg Oncol. 2015;22(11):3489–3496. doi:10.1245/s10434-015-4420-8

28. Zamzam Y, Abdelmonem Zamzam Y, Aboalsoud M, Harras H. The Utility of SOX2 and AGR2 Biomarkers as Early Predictors of Tamoxifen Resistance in ER-Positive Breast Cancer Patients. Int J Surg Oncol. 2021;2021:9947540. doi:10.1155/2021/9947540

29. Li S, Jiang F, Chen F, Deng Y, Pan X. Effect of m6A methyltransferase METTL3 -mediated MALAT1/E2F1/AGR2 axis on adriamycin resistance in breast cancer. J Biochem Mol Toxicol. 2022;36(1):e22922. doi:10.1002/jbt.22922

30. Li JJ, Wang S, Guan ZN, Zhang JX, Zhan RX, Zhu JL. Anterior Gradient 2 is a Significant Prognostic Biomarker in Bone Metastasis of Breast Cancer. Pathol Oncol Res. 2022;28:1610538. doi:10.3389/pore.2022.1610538

