## Supplementary Information for "AGR2, a potential prognostic and predictive biomarker and therapeutic to overcome drug resistance in HER2-overexpressing cancer"

Supporting Information

**S1 Material and Methods**

**Quantitative LC-MS/MS analysis**

The peptides were separated on a reversed-phase C18 Aurora column (25cm x 75μm ID, C18, 1.6 μm; IonOpticks, Australia) at a constant flow rate of 250 nL/min and an increasing acetonitrile gradient. Mobile phases were 0.1% (v/v) formic acid in water (phase A) and 0.1% (v/v) formic acid in acetonitrile (phase B). The peptides were separated by a gradient starting from 2% of mobile phase B and increased linearly to 32% for 60 min. This was stepped up to 95% of mobile phase B where it was maintained for 7 min. The injection volume was 5 μL, equivalent to a loading of 200 ng per sample.

The timsTOF Pro mass spectrometer was operated in positive ion polarity with TIMS (Trapped Ion Mobility Spectrometry), and PASEF (Parallel Accumulation Serial Fragmentation) modes enabled. The accumulation and ramp times for the TIMS were both set to 100 msec., with an ion mobility (1/k0) range from 0.62 to 1.46 Vs/cm. Spectra were recorded in the mass range from 100 to 1,700 m/z. The precursor MS Intensity Threshold was set to 2,500 and the precursor Target Intensity set to 20,000. Each PASEF cycle consisted of one MS ramp for precursor detection followed by 10 PASEF MS/MS ramps, with a total cycle time of 1.16 s.

**Supplemental Data**

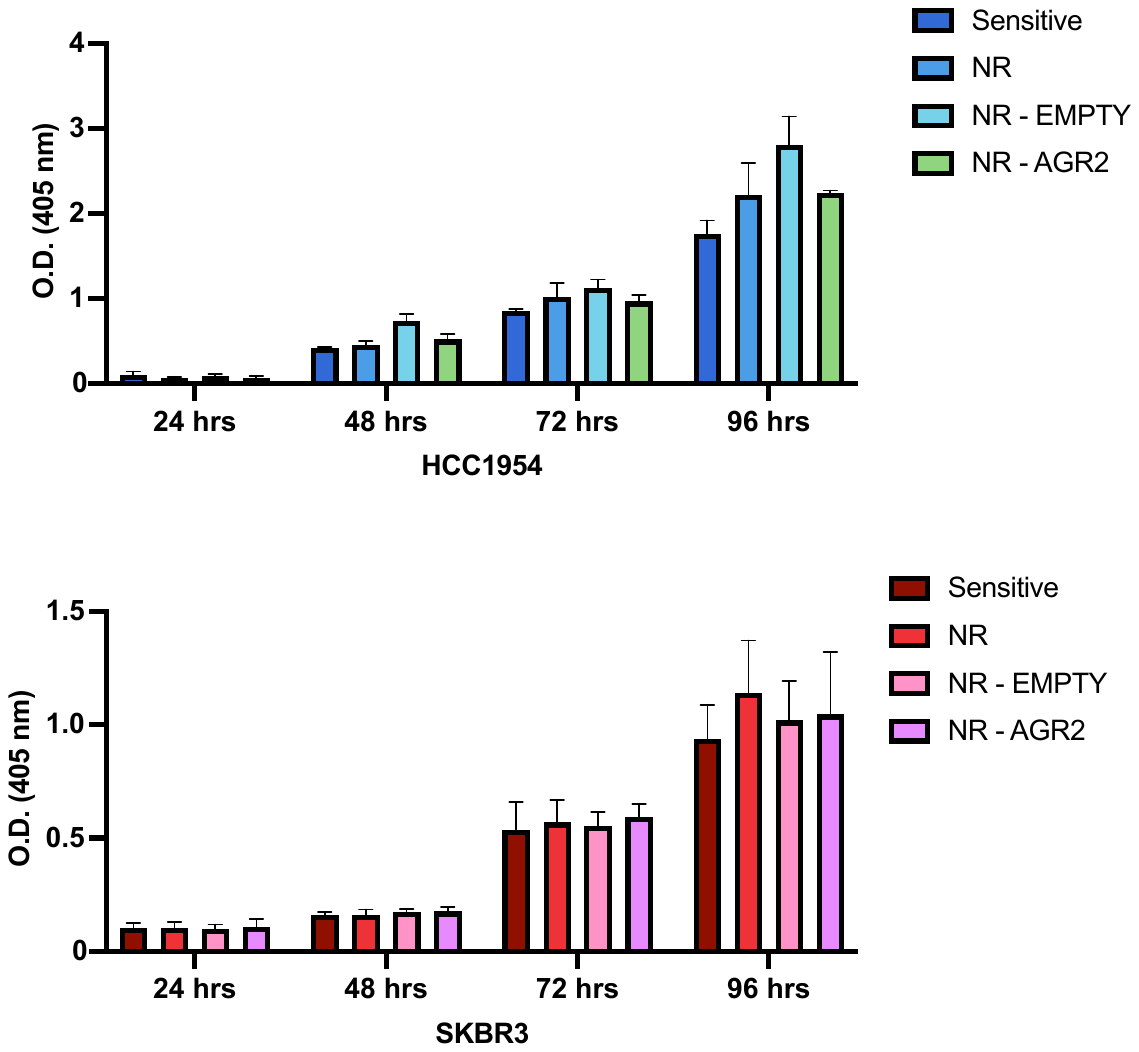
 **Figure S1. Ectopic expression of AGR2 does not affect proliferation rate of neratinib-resistant cell variants.** Acid phosphatase assay results at 24, 48, 72, and 96 hrs. The data represent the means from three independent experiments, with error bars showing the SEM of the means. OD_405_, optical density at 405nm.

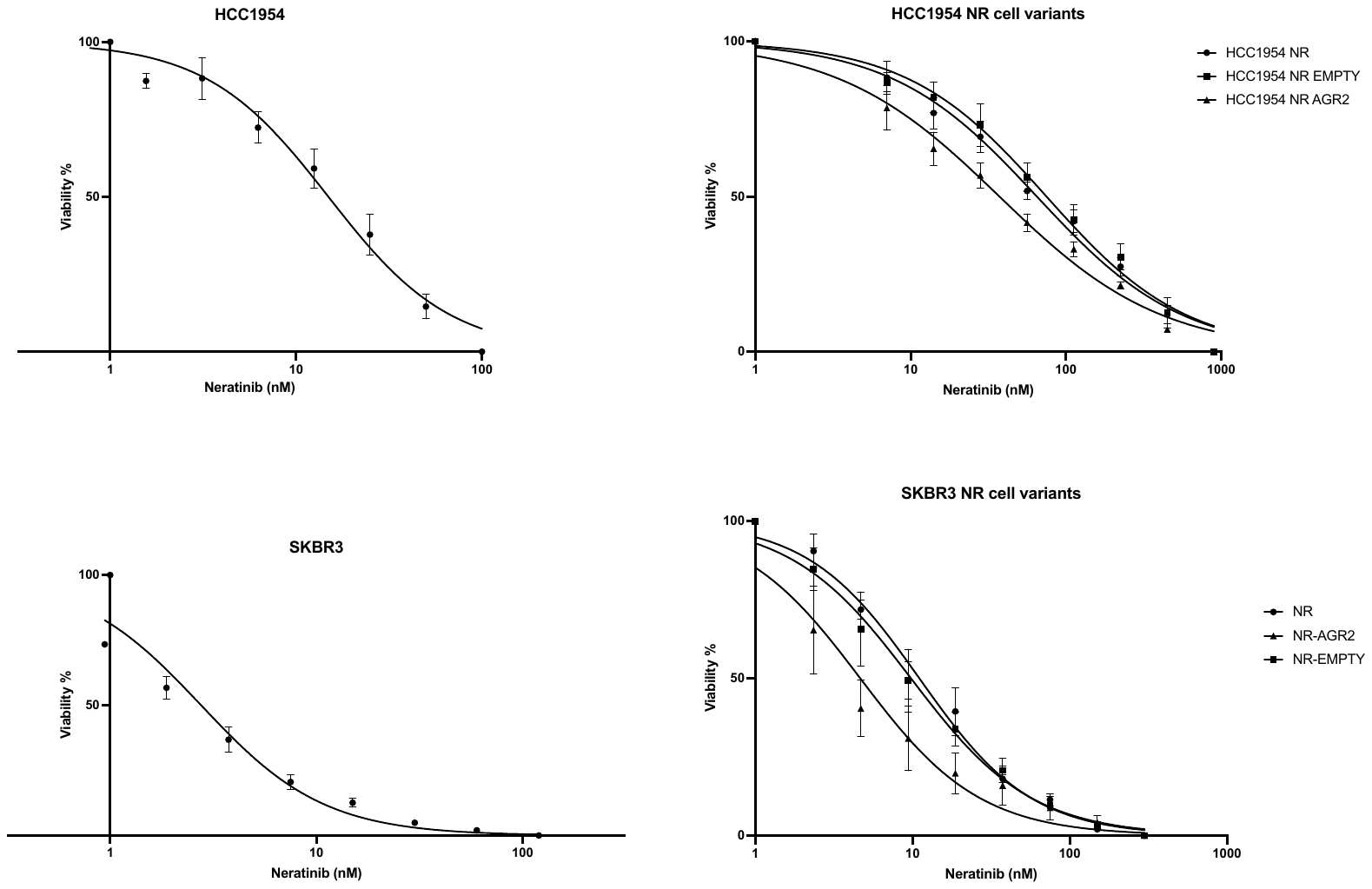
**Figure S2. Drug sensitivity assay–dose response curves of HCC1954 and SKBR3 cell variants.** Cytotoxicity assay on HCC1954 and SKBR3 cell variants after 5 days of neratinib treatment. IC_50_ curves represent *n*=3 biological repeats as mean ± SEM.
